# A Large-Scale Concordance Study of Toxicity Findings Across Preclinical Species and Humans for Small Molecules and Biologics in Drug Development

**DOI:** 10.64898/2026.01.29.702667

**Authors:** Xin Liu, Fan Fan

## Abstract

Translating preclinical safety findings into reliable insights for human risk assessment remains a fundamental challenge in drug development. Prior preclinical-clinical concordance studies have been constrained by limited drug coverage, reliance on identical-term matching for adverse events (AEs), and insufficient consideration of species, modality, exposure, and biological or mechanistic context. To address these gaps, we assembled a large cross-species concordance dataset, integrating standardized preclinical and clinical safety data for 7,565 marketed and investigational drugs from PharmaPendium and OFF-X. Our framework employs likelihood ratios to reduce prevalence bias and extends concordance assessment beyond identical-term matches to include semantically and mechanistically related AE pairs. Stratified analyses by species, modality, and exposure-matched subsets further refined translational relevance, while integration of on- and off-target annotations supports mechanistic interpretation and potential screening. Using this approach, we identified 850 significant identical-term AEs and 2,833 additional unique endpoints from cross-term associations. To promote reproducibility and transparency in animal research, we provide open access to the analytic code and statistical results via an interactive web application. An accompanying multi-agent AI system (ToxAgents) enables standardized querying and interpretation of concordance results. Together, these resources extend previous foundational efforts and establish a shared, data-driven platform to advance translational safety science, support evidence-based study design aligned with the 3Rs, and ultimately contribute to the development of safer medicines to improve human health.

## INTRODUCTION

Advancing drug safety assessment relies on bridging the gap between animal toxicology findings and clinical outcomes, a challenge with far-reaching consequences for R&D efficiency, regulatory success and patient safety *(Waring et al., 2015; Amorim et al., 2024)*. Over the past two decades, numerous concordance studies have quantified cross-species translation across endpoints and datasets (Olson et al., 2000; Tamaki et al., 2013; Clark, 2015; Monticello et al., 2017; Clark and Steger-Hartmann, 2018; Giblin et al., 2021), as summarized in **Supplementary Table 1**.

Early work by Olson et al. (Olson et al., 2000) within the International Life Sciences Institute’s Health and Environmental Sciences Institute (ILSI-HESI) quantified cross-species sensitivity by analyzing 150 drugs. They reported high concordance (>80%) for hematological, gastrointestinal, and cardiovascular toxicities, whereas cutaneous toxicities were the least concordant (<35%). Tamaki *et al*. (Tamaki et al., 2013) similarly examined 142 approved drugs in Japan using sensitivity, finding that infection, hematological, ocular, and injection-site reactions reached approximately 70% concordance, while cardiovascular, neurological, and cutaneous toxicities remained low (<30%). Monticello et al. (Monticello et al., 2017), from The International Consortium for Innovation and Quality (IQ), evaluated predictive values for 182 drugs, reporting a positive predictive value (PPV) of 43% and a negative predictive value (NPV) of 86%, indicating that the absence of toxicity in animal studies strongly predicts the absence of toxicity in the clinic. For non-human primates, predictive values were higher for nervous system and gastrointestinal toxicities.

Advances in data curation over the past decade, exemplified by resources such as PharmaPendium (PP), which compiles FDA and EMA regulatory documents, reference books such as the Meyler’s Side Effects of Drugs, and biomedical literature, have facilitated concordance analyses at larger scale and granularity (ranging between 2,259–3,815 drugs). Clark et al. (Clark, 2015; Clark and Steger-Hartmann, 2018) and Giblin et al. (Giblin et al., 2021) applied likelihood ratios to identify endpoints such as arrhythmia, QT prolongation, and drug-specific antibodies responses with strong translational value.

These larger-scale approaches have revealed important cross-species relationships; however, many prior studies have focused predominantly on approved drugs, introducing “survivor’s bias” as these drugs’ adverse events (AEs) are less severe or clinically manageable *(Clark and Steger-Hartmann, 2018)*. Moreover, systematic stratification by drug modality and animal species has been often overlooked, thereby hindering focused concordance analysis of the distinct toxicity profiles of small molecules (SM) and biologics (Bio). The absence of pharmacokinetic exposure control further confounds interpretation, as toxicities observed in animal models, often at high doses, may not accurately represent risks relevant to clinical settings with relevant efficacious doses *(Van Norman, 2019)*. Therefore, ensuring that concordance measures are based on comparable exposure levels across species helps make them more predictive of clinically relevant toxicities. Analytical inconsistencies also arise from the use of different statistical frameworks such as sensitivity, specificity, predictive values, or likelihood ratios, which can yield divergent conclusions *(Monticello et al., 2017; Bailey and Balls, 2019a; Giblin et al., 2021)*. Reliance on strict identical-term mappings can obscure semantically or mechanistically related phenotypic endpoints, thereby limiting the detection of translational signals. Mechanistic understanding is also constrained by limited integration with on- and off-target or biological pathways linked to associated AEs. Finally, restricted access to proprietary datasets and analytic results impedes application by the broader scientific, regulatory and pharmaceutical communities. These gaps outlined clear opportunities to advance concordance methodology. We address these needs through a robust, data-rich framework for cross-species comparison that enhances translational assessment, supports more rational species selection, and advances the “Reduction” and “Refinement” principles of the 3Rs in animal research by reducing redundant or low-yield animal studies and optimizing study design towards endpoints with higher expected human relevance (Törnqvist et al., 2014).

In this context and building upon the prior efforts, our study introduces several key methodologically substantive improvements (**Fig. 1**). First, we assembled a large dataset by integrating PharmaPendium with OFF-X, enabling standardized analysis of preclinical and clinical safety data across investigational and approved drugs. Animal and human safety endpoints harmonized using MedDRA were used to support consistent cross-species analyses across five major preclinical species (mouse, rat, rabbit, dog, monkey). Second, our analytic strategy prioritizes likelihood ratios to minimize prevalence bias and systematically extends beyond identical-term mapping to include semantically and mechanistically related, non-identical AE pairs. This design captures a broader and more biologically realistic spectrum of translational relationships. Third, stratified analyses by species, modality, and pharmacokinetic exposure control provide context-dependent insight into when animal data most meaningfully inform human risk. Fourth, integration of systematic on- and off-target annotation, together with pathway analysis, links observed concordance patterns to plausible molecular mechanisms. Finally, we provide access to the concordance results through an interactive web platform and an AI-powered multi-agent system, enabling users to query and explore results in in a standardized and reproducible manner. By releasing the code and complete outputs in a standardized, accessible format, our framework enables drug developers, regulatory authorities, and the broader scientific community to explore and harness the power of concordance findings in real-world decision-making contexts. These openly shared scientific resources directly support reproducibility in animal research (Frommlet, 2020). Collectively, we intend this platform to serve as a durable foundation for continued methodological refinement and innovation, aligned with our goal of delivering safer therapeutics and improved patient outcomes.

**Fig. 1.**
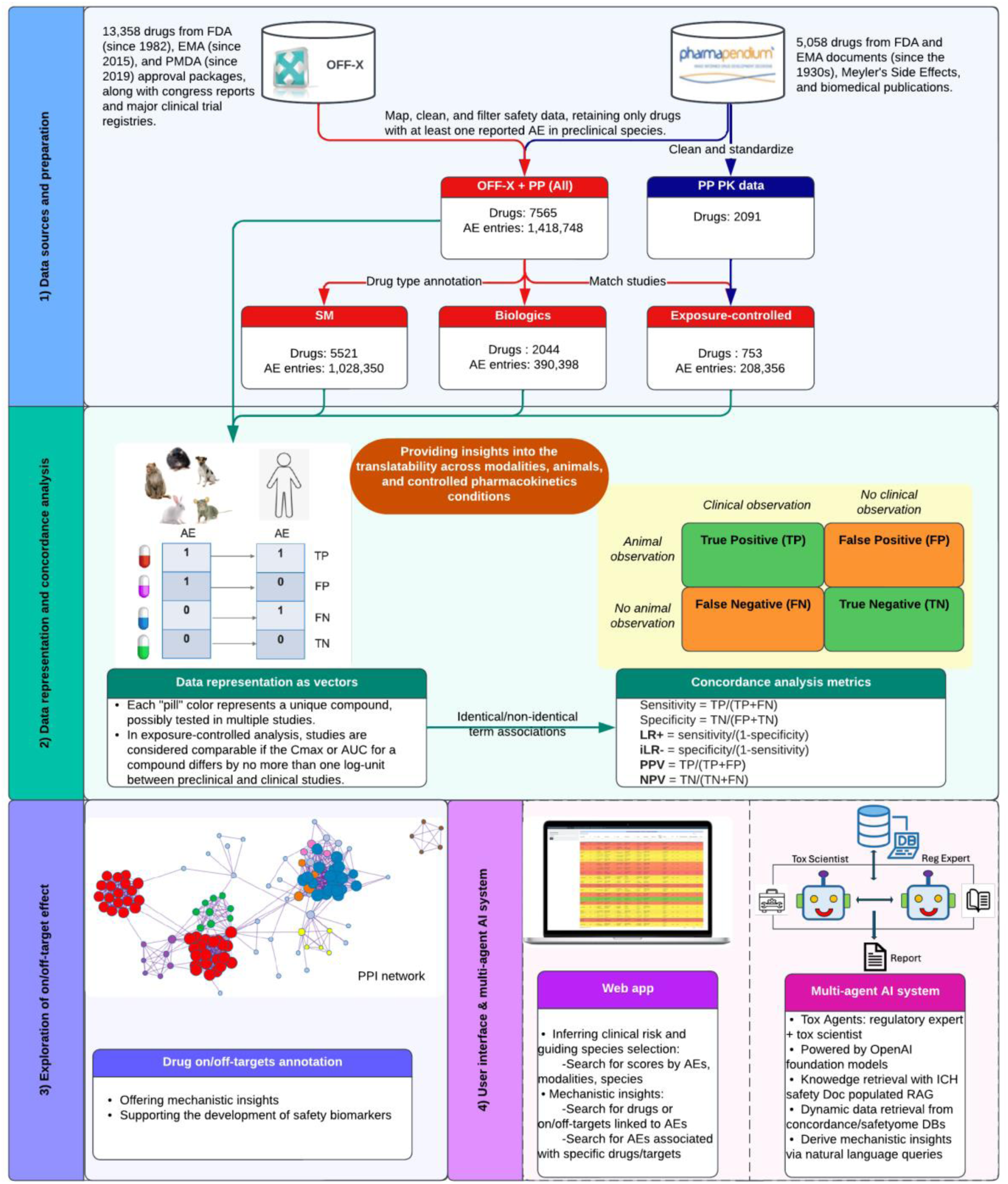
Overview of data preparation and analysis workflow. AE data encoded in MedDRA terms are sourced from PharmaPendium and OFF-X API. Drugs are consolidated and categorized by modality, and pharmacokinetic (PK) data are retrieved for exposure-controlled analysis. Preclinical toxicity studies focus on mouse, rat, rabbit, dog, and monkey. Concordance is evaluated using Fisher’s Exact Test, with likelihood ratios (LR+, iLR−) and predictive values (PPV, NPV) assessing AE predictiveness. Drug targets/off-targets, pathways, and PPI networks are explored to provide mechanistic insights and to prioritize potential safety targets for further evaluation. Findings are made accessible through a web app and a multi-agent AI system for safety assessments.

## MATERIALS AND METHODS

### Dataset and preparation

AE data from both preclinical and clinical studies were compiled from two licensed databases: Elsevier’s PharmaPendium (https://www.pharmapendium.com, accessed September 2024) and OFF-X (https://targetsafety.info, accessed November 2024). Both databases contain preclinical toxicology findings and clinical safety observations for the drugs indexed. In each source, AEs are coded using the Medical Dictionary for Regulatory Activities (MedDRA) terminology. From PharmaPendium, extracted fields included drug metadata, MedDRA mappings for each reported AE, and alert-related information (e.g., study phase, study details, species). Where available, pharmacokinetic (PK) measures such as dose, route of administration, maximum observed plasma concentration (Cmax), and area under the concentration–time curve (AUC) were also obtained, enabling quantitative exposure comparisons between preclinical and clinical datasets. From OFF-X, safety records retrieved via the API, noting that PK measures are not available in this source. For both datasets, post-marketing events were excluded because such reports are subject to less stringent criteria and variable reporting conditions (Clark and Steger-Hartmann, 2018). In OFF-X, alerts labelled “refuted/not associated” were excluded to remove disproven associations. A complete list of extracted variables, along with example data files and the Python code used for data processing and analysis, is available in the project’s GitHub repository.

Drug identifiers varied across the source datasets; therefore, we implemented a hierarchical matching strategy to ensure consistent compound alignment. Harmonized main drug names were used as the primary matching method, followed by drug synonyms from the source datasets as a secondary matching strategy. If no match was identified by name or synonym, additional identifiers were used sequentially, including PubChem Compound ID (CID), Chemical Abstracts Service (CAS) numbers, and Simplified Molecular Input Line Entry System (SMILES) notation for chemical structure representation. CID, CAS, and SMILES data were sourced or validated using DrugBank (https://go.drugbank.com/, accessed October 2024) and PubChem where not directly available from the source files. For drugs successfully matched across sources, the OFF-X “drug main name” was retained as the unique identifier in the final dataset. For PharmaPendium drugs without an OFF-X match, the original PharmaPendium name was preserved. This approach ensured a single, consistent identifier per drug entity and prevented duplication across sources. Each drug was additionally assigned with modality categories (small molecule or biologics), based on OFF-X metadata or DrugBank product annotations.

For preclinical data, five major preclinical species (mouse, rat, rabbit, dog, and monkey) were included. Species names were normalized to enable species-stratified concordance calculations. In total, 13,949 unique AE terms were retrieved from the two databases, with a detailed species and modality level breakdown presented in **Supplementary Figure 1**.

The combined dataset was further filtered to include only drugs with at least one AE reported in both preclinical and clinical studies, resulting in a total of 7,565 drugs. This step ensured relevance for evaluating translatability between preclinical and clinical phases. Duplicate records of the same drug-species-AE combination were removed, retaining only one instance to avoid bias weighting of repeated observations from datasets. For exposure-controlled analysis, compound exposure was quantified using normalized Cmax (μg/mL) or AUC (μg·h/mL). Animal and human studies were deemed comparable if the exposure difference was within one log-unit, indicated by |Δlog10Cmax| or |Δlog10AUC| ≤ 1 *(Wright et al., 2023)*.

The resulting harmonized dataset contained AE records (phenotype and organ systems) for each drug, stratified by species (as well as an “all-species” aggregate), and modality (small molecule, biologic, or all modalities combined). These groupings served as the basis for concordance analysis summarized in **Supplementary Table 2**.

### Data representation and concordance assessment

We evaluated the concordance between preclinical and clinical AE observations by constructing 2×2 contingency tables for each species-observation pair, using findings from animal studies as predictors for clinical outcomes. Compounds were classified into four categories based on the presence or absence of AEs in both studies. True positive (TP; plural, TPs) refers to an AE observed in both the preclinical species and humans. False positive (FP; plural, FPs) is an AE observed in preclinical species but not in humans, while false negative (FN; plural, FNs) describes an AE observed in humans but not in preclinical species. A true negative (TN; plural, TNs) is an AE absent in both. Note that true negative counts are estimations, as not all possible observations were measured or reported for every compound.

To determine the statistical significance of these associations, we employed two-tailed Fisher’s Exact Test with a Bonferroni correction within each system organ class (SOC; plural, SOCs) to adjust for multiple comparisons, thereby reducing the risk of false-positive results. Associations were considered significant if the adjusted p-value was <0.05. To broaden the predictive utility of available preclinical endpoints, we evaluated association and concordance not only for identical AE matches but also for non-identical endpoints within the same SOC. This approach enables the detection of semantically or mechanistically related endpoints that may be overlooked by identical-term mapping alone.

To quantify translatability, we primarily focused on likelihood ratios. To maintain inclusivity for this analysis, phenotypes with zero FPs were not excluded from the analysis. For example, gastrointestinal perforation in dogs exhibited 127 TPs and LR+ of infinity due to zero FPs. While such cases yield extreme LR estimates, their inclusion avoids arbitrary exclusion of potentially informative phenotypes and preserve transparency for downstream interpretation. However, caution is warranted when interpreting phenotypes with small TP counts and infinite LR+, as these estimates may be unstable. To complement LRs, we additionally reported PPV and NPV, which reflect the probability that a clinical adverse event will be present or absent in response to a drug, given preclinical findings. Since predictive values are affected by both the prevalence of the AE and the model’s predictive accuracy, they offer practical utility for assessing post-test clinical risk. All metrics were computed as illustrated in **Fig. 1**, and our findings will evolve as the underlying databases are updated.

To address the semantic similarities between preclinical and clinical AE terms, we utilized the OpenAI text-embedding-3 model to generate dense vector representations for all phenotype terms. By grouping preferred term (PT; plural, PTs) based on SOC, we ensured that comparisons were made within biologically relevant categories. Pairwise cosine similarity scores were then calculated between each preclinical and clinical PT, allowing us to quantify semantic relatedness. All embeddings and similarity computations were performed programmatically through the OpenAI API and standard scikit-learn functions in Python.

### Retrieving drug targets linked to toxicity in preclinical and clinical AEs

To explore the potential biological mechanisms underlying toxicity, on- and off-targets for each consolidated drug were retrieved from DrugBank and the OFF-X API. These data were used to annotate drugs in different categories for each associated AE.

Pathway and functional enrichment analyses were conducted in R with gprofiler (Kolberg and Raudvere, 2025), employing the DisGeNET database (Piñero et al., 2017) to explore relevant biological pathways and processes. Multiple testing correction was applied using the Benjamini-Hochberg (BH), with significance defined as an adjusted p–value < 0.01. Protein-protein interaction (PPI) analysis was performed using the STRING database (Szklarczyk et al., 2023) to investigate potential interactions among targets. The network was visualized using RCy3 *(Gustavsen et al., 2019)*, an R interface for Cytoscape, to facilitate the interpretation of complex interactions. To refine the network and focus on key interactions, the MCODE algorithm *(Bader and Hogue, 2003)* was applied to identify densely connected sub-networks, which were further analyzed for functional enrichment to determine their biological roles and contributions to toxicity. This analysis highlighted potential key proteins and pathways driving the observed adverse effects, providing valuable mechanistic insights and testable hypotheses to understand drug-induced toxicity.

### Web application

A web app was developed using the Shiny framework in R to provide a user-friendly platform for exploring AE concordance data. This tool enables detailed searches for AE translatability scores by modality and species, supporting evaluation of animal-to-human translation and informed species selection in preclinical studies. Users can also search drugs and their on- or off-targets associated with specific AEs, or identify AEs linked to particular drugs or targets, thereby facilitating mechanistic insights into potential safety.

To enhance accessibility and interpretation of concordance results, we further developed an AI-powered multi-agent system, ToxAgents, for standardized interpretation, and real-world safety evaluation. The generative component is powered by OpenAI’s GPT-4.1 model, while the text-embedding-3-small is used to construct the retrieval-augmented generation (RAG) system. This RAG system employs CrewAI’s default process for knowledge retrieval, which includes chunking, embedding, and storing ICH safety documents in a vector database, with the same embedding used for subsequent retrieval. In this context, the Investigative Toxicity Crew is composed of two agents: the regulatory affairs expert agent, which leverages a RAG system consisting of 26 ICH safety documents as its knowledge base, and the investigative toxicology scientist agent, who accesses the concordance and safetyome tools (Liu et al., 2021).

The safetyome and concordance data are preloaded into separate PostgreSQL databases. Both the safetyome and concordance tools make extensive use of LLMs to generate SQL queries tailored to user inputs, thereby fetching the most relevant data. This retrieved data is then incorporated into the agents’ reasoning processes. Finally, two sequential tasks are designed to ensure that the agents evaluate user queries from both regulatory and scientific perspectives.

For sanity check, a QA list comprised of preclinical safety-related questions, and their expected answers is provided. This list simulates the scenarios and real-world questions encountered in daily work. The goal of the sanity check system is to compare the agent-generated answers with the expected responses. To accomplish this, the system is heuristically constructed by combining the output of the multi-AI agent system with the expected human answer into a single prompt. A native LLM then acts as a judge, evaluating semantic similarity between them. For the case studies, we also assessed reproducibility by running each question five times and measuring the consistency of the generated outputs. In addition, we evaluated robustness to prompt variation by comparing the outputs from the original prompt for each case to those generated from five paraphrased versions, quantifying the stability of the responses under varied phrasings.

The ToxAgents implementation code and installation scripts are open source and available on the ToxAgents GitHub repository.

## RESULTS AND DISCUSSIONS

### SOC-based AE Frequency and Concordance Analysis

We assessed AE distribution at the SOC level across therapeutic modalities and preclinical species. As shown in **Fig. 2** (Panels A and B), “investigations”, primarily representing laboratory test abnormalities, emerged as the predominant AE category in both human (SM 11.6%; Bio 9.5%) and animal studies (SM 30.9%; Bio 27.3%). “General disorders and administration site conditions” also exhibited high frequencies, with biologics showing slightly higher relative percentages than small molecules (Bio 9.4% vs. SM 7.9% in humans; Bio 10.8% vs. SM 8.1% in animals), which likely reflects parenteral administration (e.g., injections, infusions) of biologics, and may result in localized reactions at the injection site.

**Fig. 2.**
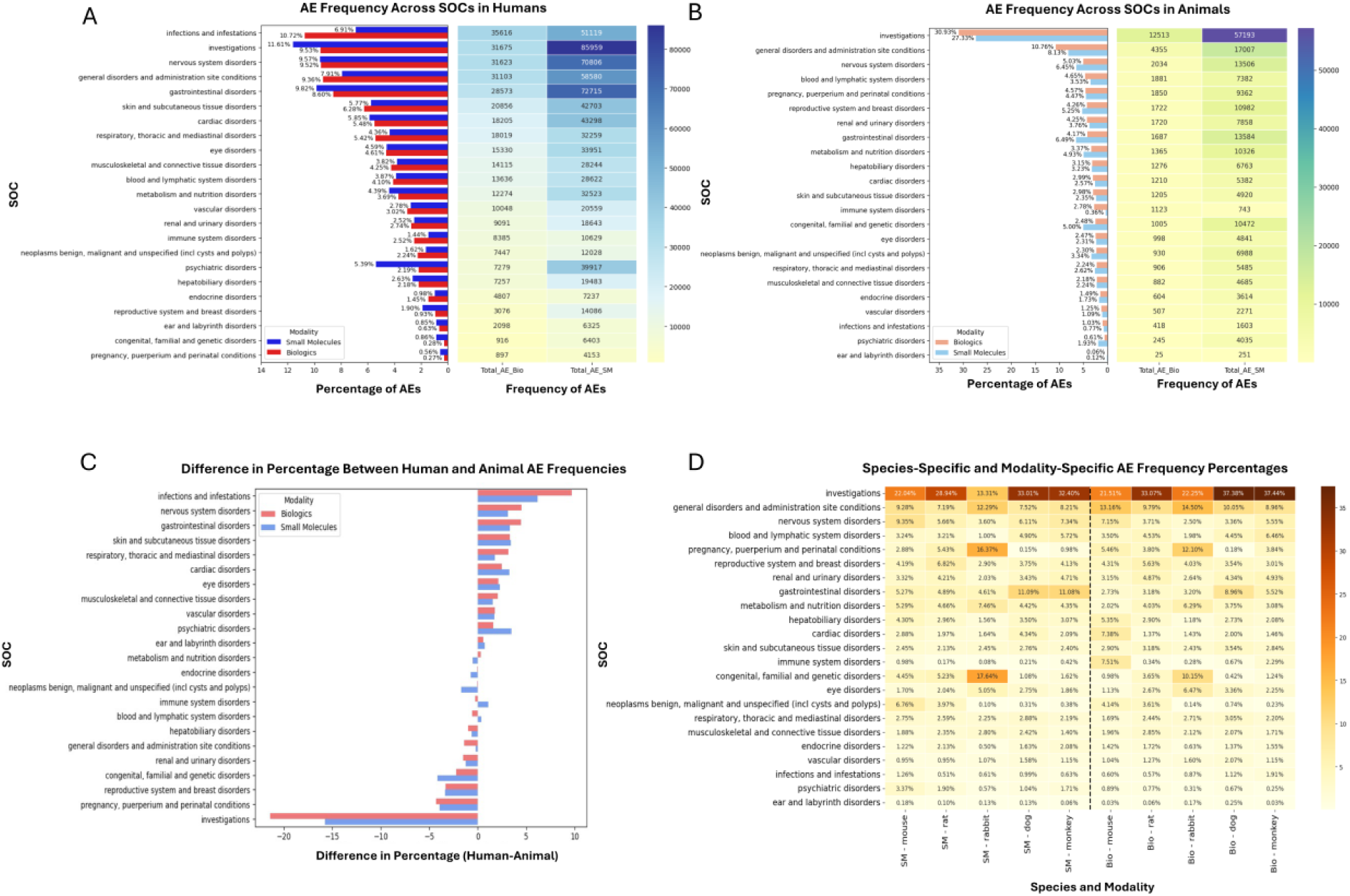
SOC-based AE frequency analysis across modalities and species. (A) Comparison of adverse event (AE) frequencies across SOCs in humans for biologics and small molecules. The bar graph indicates percentage distributions, and the heatmap visualizes absolute AE frequencies. (B) Corresponding AE distributions from preclinical studies. Bar graph and heatmap demonstrate relative and absolute AE occurrences, respectively, across modalities. (C) Difference in percentage between human and animal AE frequencies (calculated as human percentage minus animal percentage) stratified by SOC and drug modality, highlighting SOCs disproportionately represented in either human or animal studies. Positive values indicate SOCs more frequently reported in human studies; negative values indicate higher reporting in animal studies. (D) Species-specific and modality-specific detailed heatmap illustrating AE frequency percentages across SOCs. Darker shading reflects higher frequencies, providing detailed insights into interspecies and modality variability.

Significant discrepancies between human and preclinical studies were observed across several categories (**Fig. 2**, panel C). “Infections and infestations”, as well as “psychiatric disorders”, were notably underrepresented in animal studies (<1%) when compared to humans (infections: Bio 10.7%, SM 6.9%; psychiatric disorders: SM 5.4%, Bio 2.2%), which may relate to differences in immune status, microbial exposure, and the scopes of behavioral assessments between animals and humans *(Dinan and Cryan, 2017; Huggins et al., 2019; Burger et al., 2023)*. Conversely, “reproductive and breast disorders”, “congenital, familial, and genetic disorders”, and “pregnancy and perinatal disorders”, were more frequent in animals, suggesting different study protocols in humans and animals due to practical and ethical considerations. Species-specific differences were also evident (**Fig.2**, panel D). For example, “gastrointestinal disorders” were more frequent in dogs and monkeys, consistent with their known susceptibility comparing to rodents *(Clark and Steger-Hartmann, 2018)*. Additionally, the disproportionately high incidence of “pregnancy and perinatal toxicities” observed in rabbits aligns with their common utilization in reproductive toxicology studies *(Foote and Carney, 2000; Moxon et al., 2023)*. In this analysis, certain SOCs, including “Investigations,” “Congenital, familial and genetic disorders,” and “General disorders,” contain phenotypes that span multiple organ systems. While some terms are clearly organ-associated, others cannot be assigned with certainty to a single organ system within the MedDRA hierarchy. Selectively reassigning or excluding these terms at this stage would reduce dataset completeness and could introduce bias into the concordance analysis. To safeguard data integrity and ensure a baseline that is directly comparable with prior concordance studies, we retained the original SOC assignments in this work. Future analyses will investigate systematic, data-driven methods, including NLP-based approaches that map directly from phenotype-level terms to organ systems, to enable consistent, reproducible, and unbiased reassignment at scale.

The translational concordance of AEs between humans and preclinical species at the SOC level was evaluated as described in the Methods section, which details the contingency table framework, statistical testing, and significance criteria used. To quantify translatability, we calculated positive likelihood ratio (LR+) and inverse negative likelihood ratio (iLR–) as illustrated in **Fig. 1**. These metrics are independent of event prevalence and incorporate both sensitivity and specificity *(Bailey et al., 2014; Bailey and Balls, 2019b; Shreffler and Huecker, 2024)*. In this study, LR+ quantifies the increased probability of an AE occurring in humans when it has been observed in preclinical species, measuring the strength of positive concordance. Conversely, iLR− reflects the reduced likelihood of an AE manifesting in humans when it has not been detected in preclinical species, representing the strength of negative concordance. Likelihood ratios range from zero to infinity, with 1 indicating no predictive power; values above 1 demonstrate increasing predictivity, and ratios exceeding 10 are generally considered strong *(Clark, 2015; Monticello et al., 2017)*. Likelihood ratios effectively quantify the predictive value of findings from animal studies. Notably, a highly statistically significant association combined with a low likelihood ratio suggests that the animal observation does not predict the clinical outcomes.

Applying this framework, we performed concordance at the SOC level as shown in **Fig. 3**. Across all modalities, “infections and infestations” exhibited the highest predictivity (LR+ >10) evident for both small molecule and biologics. This may reflect similar immunological mechanisms between non-human primates and humans and the effect of immunomodulatory agents. This was followed by “immune system disorders”, “renal and urinary disorders”, and “musculoskeletal and connective tissue disorders”, although these showed only moderate predictive strength (LR+ < 5). Predictivity of biologics for “immune system disorders” was higher than that observed for small molecules, consistent with the known propensity of biologics to elicit immune-mediated adverse reactions. Such toxicities include cytokine release syndrome driven by T-cell activation, as well as immune-mediated organ injury such as checkpoint inhibitor-associated hepatitis, which have been qualitatively recapitulated in preclinical species, particularly in non-human primates and humanized mouse models (Sauna et al., 2018; Yong et al., 2020; Llewellyn et al., 2021; Kroenke et al., 2024). SOCs such as “nervous system disorders”, “psychiatric disorders”, “investigations”, “general disorders” and “pregnancy and perinatal conditions” showed low LR+ values, indicating weaker translational predictivity. Panel B shows a breakdown by species. While median LR+ values were broadly similar across most species, monkeys demonstrated higher LR+ medians (SM median = 4.18, Bio median = 3.74) compared with other species, suggesting stronger predictivity. Panel C stratifies LR+ by SOC for each modality and species. For example, in “infections and infestations”, monkeys showed the highest predictivity for both modalities, along with strong signals for “immune system disorders”, “musculoskeletal disorders”, “endocrine disorders”, and “blood and lymphatic system disorders”. Dogs displayed high predictivity for “gastrointestinal disorders” and “vascular disorders”, whereas rabbits performed relatively well for “pregnancy conditions” and “eye disorders” in small molecules. Mice showed relatively higher predictivity for “congenital and genetic disorders”. However, it should be noted that, when aggregating at the SOC level, most LR+ values fell within the low to moderate range. This finding is consistent with previous work showing that, although SOC-level relationships are often highly statistically significant, likelihood ratios may not indicate strong predictivity (Clark and Steger-Hartmann, 2018). In that study, the highest predictivity was observed for infections and infestations (LR+ = 9.6), while general disorders, investigations, and pregnancy-related conditions were among the lowest (LR+ < 3), broadly aligning with the pattern seen in our results.

**Fig. 3.**
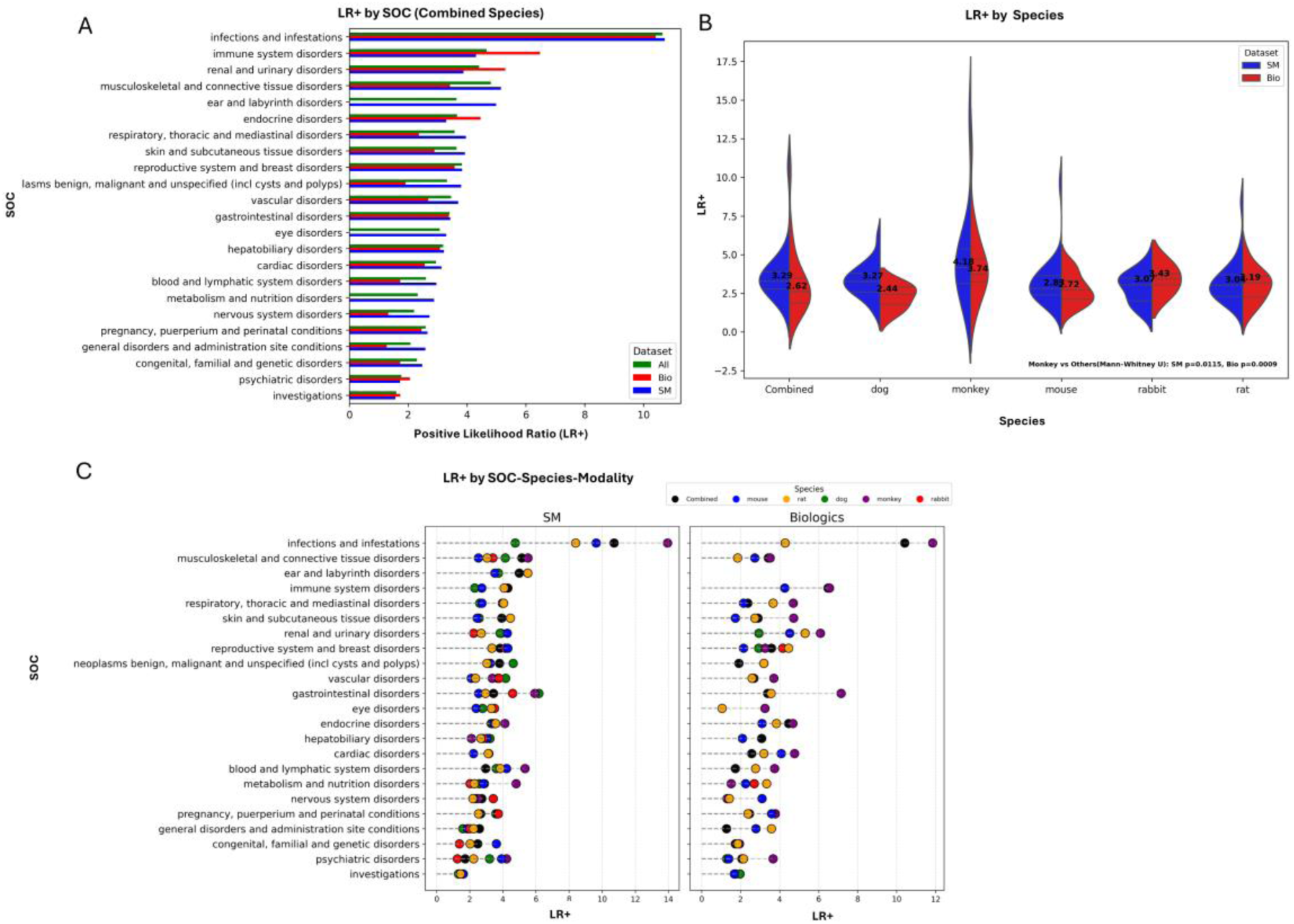
Positive likelihood ratios at SOC level by species and modality. (A) LR+ values for combined species across SOCs. Bars depict LR+ for all modalities (green), small molecules (blue) and biologics (red). (B) Violin plots of LR+ distributions by species, comparing SM (blue) and Bio (red). Black dashed lines mark medians, with values displayed within each plot. monkeys show notably higher LR+ compared with other species. (C) LR+ values for individual SOCs stratified by modality and species. Each point represents LR+ for a specific SOC–species pair in SM or Bio, with colors indicating species.

Regarding negative predictivity, the iLR– values indicated that preclinical models have a limited ability to predict the absence of human AEs (**Supplementary Table 3**). Across both modalities, most iLR– values were close to 1, indicating little evidence that the absence of an AE in preclinical studies meaningfully reduces the probability of observing the corresponding event in humans. This result is consistent with previous reports (Bailey et al., 2014; Clark, 2015; Clark and Steger-Hartmann, 2018).

While SOC-level analysis is useful for identifying broad patterns of translational concordance across organ systems, most SOCs exhibited moderate to low predictive values. This limited predictive strength may reflect the aggregation of a broad spectrum of heterogeneous AEs with diverse biological mechanisms and varying predictive potentials. Accordingly, PT-level analysis provides a greater resolution for phenotype-specific translational assessment.

### Phenotype-Level Concordance Using Matched Terms

Building on the SOC-level findings, we next conducted a detailed PT-level concordance analysis, stratified by drug modality and including the subset with exposure-controlled data to minimize pharmacokinetic confounding.

**Fig. 4**, panel A highlights the overlaps among statistically significant AEs observed across stratified datasets. Across all modalities, 850 significant phenotypes were identified, including 780 for small molecules, 237 for biologics, and 94 in the exposure-controlled subset. Although fewer significant PTs emerged from the exposure-controlled analysis, these represent a rigorous subset, offering pharmacologically precise translational concordance assesment.

**Fig. 4.**
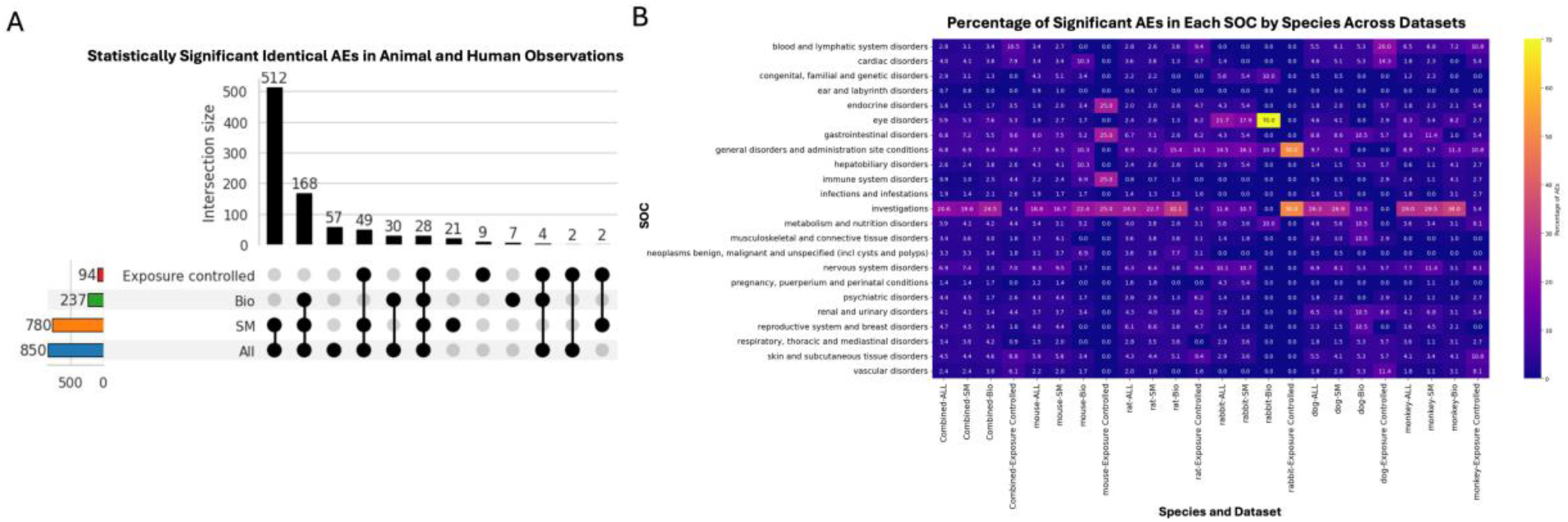
Concordance analysis at PT level between identical preclinical and clinical AEs. (A) (A) UpSet plot illustrating the overlap of statistically significant AEs between animal and human. Bar heights represent the number of significant PT terms for each dataset: All modalities (blue), Small Molecule (orange), Biologics (green), along with the Exposure-Controlled subset (red). Connected black dots indicate intersections between datasets, with the bar above each set representing the count of significant PTs shared across those datasets. (B) Heatmap displaying the percentage of statistically significant PT-level AEs identified in each SOC, across species, and modalities. Brighter colors indicate higher counts of significantly concordant terms.

The percentage-based heatmap (panel B) illustrates the distribution of statistically significant PT-level AEs within each SOC, across species and datasets. The SOC “investigations” (standard laboratory tests) consistently exhibited high proportions of significant AEs across nearly all species and modalities. Another particularly noteworthy observation was the high percentage of significant AEs within the SOC “eye disorders” in rabbits. This enrichment highlights the rabbit’s known ocular sensitivity and its established role as a preclinical model for ocular toxicity assessment, especially in biologic drug development *(21–23)*. Rabbits also showed elevated percentages of significant AEs in “general disorders and administration site conditions”, reflecting their role in evaluating local administration-related effects and associated systemic responses *(Clark and Steger-Hartmann, 2018)*. In the exposure-controlled dataset, dogs exhibited notable percentages of significant AEs in “blood and lymphatic system disorders” (20.0%), “cardiac disorders” (14.3%) and “vascular disorders” (11.4%), reflecting their known sensitivity to hematologic and cardiovascular toxicities and underscoring their utility in evaluating related safety risks in preclinical studies *(Clark and Steger-Hartmann, 2018; Koshman et al., 2024)*. Additionally, gastrointestinal disorders showed relatively high AE percentages in both dogs and monkeys compared to other species, also highlighting species specific susceptibilities *(Al-Saffar et al., 2015; Dalgaard, 2015)*.

To fully understand predictive value, we next examined individual AEs and their corresponding likelihood ratios in detail. **Supplementary Table 4** summarizes the concordance of the 25 most prevalent AEs in humans. Prevalence refers to the number of unique drugs associated with a given AE in humans, based on non-redundant AE reports. Among small molecules, several hematological and gastrointestinal AEs emerged as particularly translatable, with high LR+ values and strong statistical significance. Events such as constipation (LR+ = 21.5), rash (20.0), neutropenia (16.1), and anemia (10.1) were consistently detected in preclinical models. For biologics, infection (116.0), rash (45.2), hyperglycemia (34.4), and neutropenia (21.8) demonstrated the strongest concordance. These AEs are frequently observed in biologics targeting immune checkpoints, or cytokines *(Rider et al., 2016; Postow et al., 2018)*. In contrast, symptoms such as vomiting, fatigue, death, and seizure had low to moderate LR+ values across modalities. Vomiting and fatigue often lack clear behavioral analogs in preclinical models *(Polson and Fuji, 2012; Horn, 2014)*, while death and seizure are strongly influenced by multifactorial causes and inter-species variability *(Wang et al., 2025)*, all of which reduces the generalizability of these endpoints.

To further explore translational patterns, we examined the top 25 AEs ranked by highest LR+ across both modalities (**Supplementary Table 5**). Among small molecules, the highest LR+ AEs included gastrointestinal perforation, *Clostridium difficile* colitis, retinal vein occlusion, and macular oedema, along with several phenotypes exhibiting infinite LR+ values, such as newborn persistent pulmonary hypertension and serous retinal detachment. These events typically involve well-defined pathologies and comparatively objective endpoints, facilitating reliable detection in animal models. For example, drug-induced gastrointestinal perforation has long been modeled in animals, especially dogs, and extensive work on NSAIDs has established reproducible frameworks linking this AE to human (Lascelles et al., 2005; Mabry et al., 2021). Similarly, *C. difficile* colitis is supported by mouse models that recapitulate key clinical and histopathologic features and enable mechanistic studies (Chen et al., 2008; Theriot et al., 2014). Ophthalmic AEs such as retinal vein occlusion and macular oedema are also amenable to preclinical modeling, where laser- or occlusion-based paradigms in rodents and larger species reproduce hallmark vascular and edema phenotypes that can be quantified by fundus imaging, and histopathology (Khayat et al., 2017). For biologics, the highest LR+ values were dominated by immune-mediated toxicities, including immune-mediated hepatitis, cytokine release syndrome, thyroiditis, and broader autoimmune disorders. For example, drug-induced cytokine release syndrome has been studies in humanized mouse model with success measured by observing clinical signs and elevated human cytokines (IL-6, IFN-γ, TNF-α) that mirror human CRS (Ye et al., 2020; Nouveau et al., 2021).

Conversely, AEs with the lowest LR+ values, regardless of modality, largely included syndromic or multifactorial events, such as fatigue, neurotoxicity, developmental delay, and broadly categorized disorders affecting the cardiac, fetal, lung, skin, and liver systems (**Supplementary Table 6**). These events often lack precise analogs in animal models, are subject to broad etiologic variability, or are under-detected due to behavioral and physiological differences across species *(Van Norman, 2019; Lu et al., 2023; Wu et al., 2023)*. For instance, fatigue is a multidimensional, highly individualized symptom experience that is influenced by disease context, treatment burden, and psychosocial factors, making it difficult to map onto conserved animal phenotypes (Mead et al., 2016). Such limitations align with broader critiques of the challenges in predicting certain human toxicities from animal studies, especially for nuanced functional and neurobehavioral endpoints (Van Norman, 2019).

We also assessed the iLR– to evaluate how reliably the absence of an AE in animal studies predicts its absence in clinical settings (**Supplementary Table 7**). The majority of AEs showed relatively low iLR– values under 3, as repeatedly reported from previous concordance analyses *(Clark and Steger-Hartmann, 2018; Giblin et al., 2021)*. A subset including gastrointestinal hypermotility, testicular atrophy, spleen disorders, and medullary thyroid cancer, demonstrated higher iLR– values (>3), suggesting stronger negative predictivity. In small molecules, testicular and pancreatic disorders as well as hyperphagia were associated with relatively high iLR–values, while in biologics, teratogenicity, immune-mediated hepatitis or dermatitis, and fetal damage showed similarly high values.

We further compared identical-AE concordance metrics from our dataset with those reported in the previous large-scale analysis by Giblin et al. (Giblin et al., 2021) using 2,259 drugs, which was the only large-scale concordance analysis with publicly accessible results. As the previous study did not stratify by modality or species, our comparison was restricted to metrics aggregated across all modalities and species. Of the 188 adverse events that were identified as significant by Giblin et al. analysis, 183 (97%) were also significant in our dataset. Across all four metrics (PPV, NPV, LR+ and iLR–), scatter plots of overlapping AEs showed strong correlations between these two bodies of work (Pearson r range = 0.61–0.88), indicating general agreement despite differences in drug coverage and data sources (**Supplementary Figure 2**).

For AEs with strong predictivity with LR+ ≥ 10 in the previous study, 70% also met this threshold in our analysis. These included “drug-specific antibody present”, “renal papillary necrosis”, “blood prolactin increased”, “endometrial hyperplasia”, “occult blood positive”, “adrenal insufficiency”, “injection site discoloration”, “skin atrophy”, “neutropenia”, “iritis”, “gastrointestinal ulcer”, and “eye hemorrhage”. In all such cases, our LR+ estimates were derived from a substantially larger number of true-positive observations, in contrast to the limited counts in the previous dataset, providing greater statistical stability and robustness. In addition, our expanded dataset enabled stratifications that were not provided by previous studies. For example, “drug-specific antibody present” was significant only for biologics and was driven primarily by findings in monkeys. It also retained high concordance when compound exposure was considered. Broader coverage in our dataset doubled the number of significant AE signals identified compared with the previous analysis, increasing statistical power and enhancing the detection of translational safety signals.

Taken together, these results confirm that our expanded dataset robustly reproduces the key findings of the earlier work with improved coverage, robustness, while offering modality and species-level resolution. These improvements underscore the value of dissecting translational concordance at the level of further defined specifics (exposure, modality, and species), enabling identification of reliable safety signals and enhancing the predictive utility of preclinical models in safety assessment.

### Hepatobiliary and Metabolic Phenotypes as an Illustrative Case

To exemplify phenotype-level translatability, hepatobiliary and metabolic AEs were selected to evaluate modality- and species-specific concordance and to obtain putative mechanistic insights (**Fig. 5**).

**Fig. 5.**
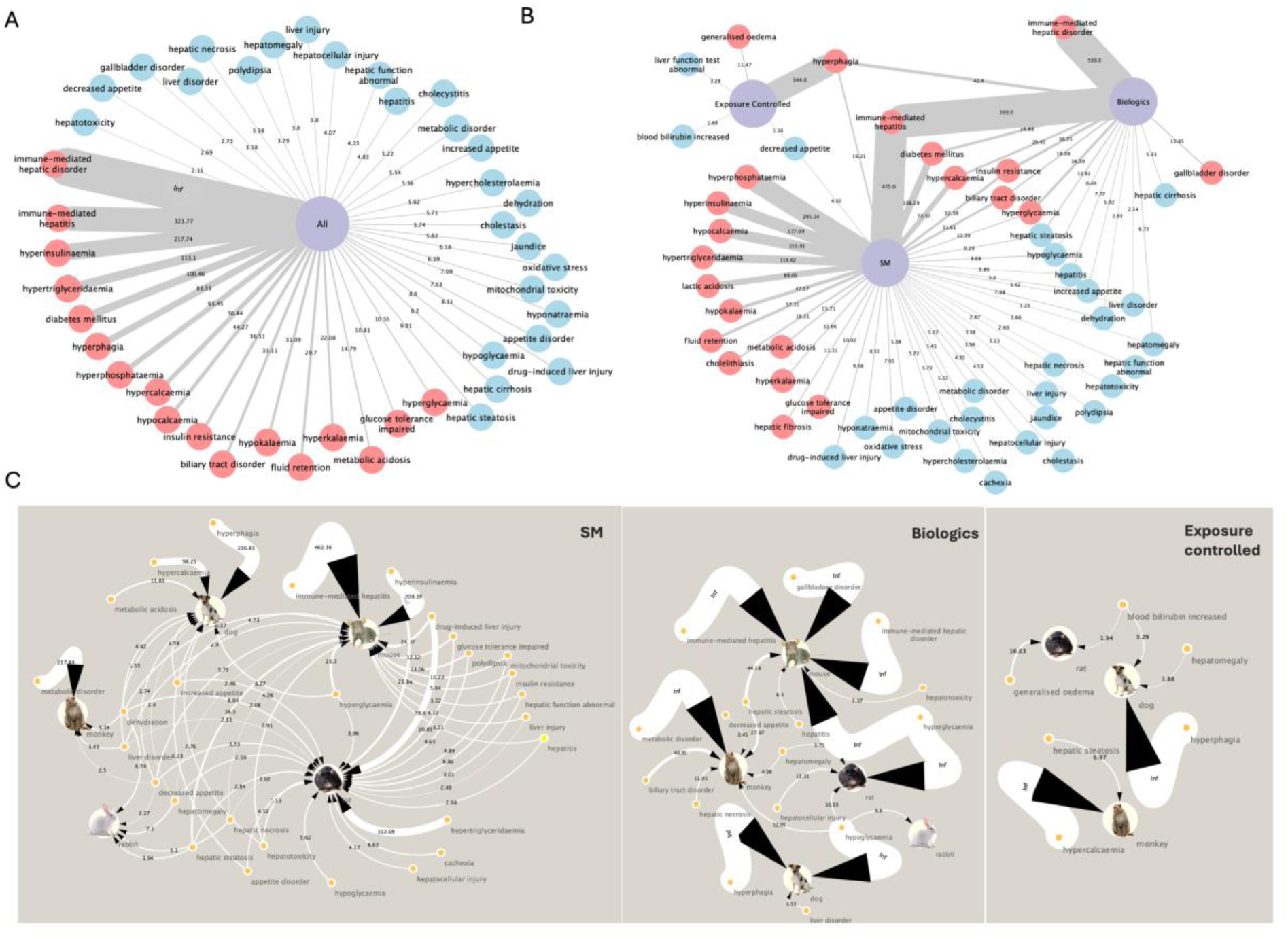
Network visualization of significant AEs within the hepatobiliary system and metabolism/nutrition disorders, based on LR+. (A) Combined analysis of all drug modalities. Nodes represent individual AEs, and edge width corresponds to LR+ values. Red and blue nodes denotes the AEs with LR+ ≥10 and < 10 respectively. (B) Stratified networks for small molecules (SM), biologics, and the exposure-controlled subset. Thicker edges denote higher LR+. (C) Species-level networks for small molecules, biologics, and the exposure-controlled subset, showing species-specific LR+ values across preclinical models. Edge width corresponds to LR+, and species icons indicate contributing data from mouse, rat, rabbit, dog, or monkey.

#### Concordance Across Modalities and Exposure-Controlled Datasets

When examining across species and without consideration of modalities, multiple hepatobiliary and metabolic AEs exhibited exceptionally high translational concordance (LR+ > 100), including immune-mediated hepatic disorder (LR+ = ∞), immune-mediated hepatitis (321.8), hyperinsulinemia (217.7), and hypertriglyceridemia (113.1). Metabolic dysregulations such as hypertriglyceridemia and hyperinsulinemia also showed high concordance. In contrast, nonspecific labels such as hepatotoxicity (LR+ = 2.2), hepatic function abnormal (4.2), or liver disorder (3.2) showed poor translational value. These terms aggregate disparate pathologies, including immune, cholestatic, and idiosyncratic liver injury, limiting their utility for mechanistic inference or preclinical model selection.

Additional delineation can be obtained when stratified by modality. SMs showed higher LR+ values for AEs associated with metabolic stress and oxidative injury, such as hyperphosphatasemia (295.1), and lactic acidosis (89.0). In contrast, biologics displayed stronger translatability for immune-mediated hepatic AEs, including immune-mediated hepatitis and hepatic inflammation. This difference likely reflects modality-specific mechanisms, where biologics more frequently trigger immune-driven responses compared to small molecules *(Leach, 2013; Shah et al., 2020)*. Notably, phenotypes such as diabetes mellitus (LR+ = 106.3 for SMs; 34.9 for biologics), insulin resistance (32.4; 36.4), and hypercalcemia (73.6; 29.5) showed robust concordance across both modalities, indicating downstream metabolic impacts are independent of mode of action. These results underscore how systemic phenotypes with convergent pathophysiology (e.g., glucose or calcium dysregulation) may serve as shared translational markers across diverse drug modalities *(Qamar et al., 2023; Jain and Lai, 2024)*.

In exposure-controlled datasets, fewer AEs met statistical significance, yet those that did, such as hyperphagia and generalized edema exhibited high LR+ values (344.0 and 11.5, respectively).

#### Species-Specific Translatability Patterns

Species-level stratification provided important insights. Mice demonstrated the strongest signal for immune-mediated hepatitis (LR+ = 462.4), reinforcing their well-established role in modeling immune–driven liver damage *(Chakraborty et al., 2015; Gerussi et al., 2021)*. Rats demonstrated high translatability for hypertriglyceridemia (112.7), in line with their known sensitivity to hepatic lipid accumulation and chronic metabolic stress *(Boivin and Deshaies, 1995; Kucera and Cervinkova, 2014)*.

Dogs exhibited broad and robust translatability for multiple AEs, including hyperphagia (230.8 in SMs; ∞ in both biologics and exposure-controlled subsets), hypercalcemia (98.2), and appetite-related disorders. Their larger body size and ability to capture behavioral endpoints (e.g., feeding behavior) enhance their sensitivity for metabolic AEs *(Martinez et al., 2020; Qu et al., 2022)*. Monkeys demonstrated high concordance for metabolic disorder (LR+ = 217.4), reflecting their physiological similarity to humans in lipid metabolism and endocrine regulation, and underscoring the relevance of non-human primates in modeling systemic metabolic responses *(Bremer et al., 2011; Friedman et al., 2017; Havel et al., 2017)*. Rabbits showed low to moderate translatability for hepatic and metabolic endpoints.

#### Mechanistic Pathway Insights via Target-Centric Annotation

To contextualize translatability in mechanistic terms, we annotated the pharmacological targets and off-targets of true-positive compounds associated with two exemplar adverse events, hyperphagia and immune-mediated hepatitis. The PPI clustering and pathway enrichment analysis were performed with the on- and off-targets (**Supplementary Figure 3**). Incorporating target-based annotation into translatability workflows enhances mechanistic interpretability by linking preclinical safety signals to underlying molecular networks.

For hyperphagia, enriched pathways included neuroactive ligand-receptor interactions, GABAergic and serotonergic signaling, monoamine GPCRs, calcium signaling, and arachidonic acid metabolism. These clusters point to CNS-mediated appetite regulation involving hypothalamic neurocircuitry and suggest that hyperphagia may reflect direct drug effects on neural and hormonal satiety control *(Morton et al., 2014; Zhuang et al., 2017; He et al., 2021; Wang et al., 2024; Qiu and Fu, 2025)*. Within these pathways, key targets can be annotated, including CNS and metabolic receptors such as HTR2A, HTR2C, HTR6 (serotonin receptors); DRD1–5 (dopamine receptors); HRH1 (histamine receptor); CHRM3 (muscarinic acetylcholine receptor); ADRA1A/B and ADRA2B/C (adrenergic receptors); NR3C1 (glucocorticoid receptor); among others. Annotating these safety targets enables focused receptor-binding and functional assays to confirm mechanisms, and guides pathway-informed monitoring and mitigation strategies. This approach links preclinical hyperphagia to actionable molecular mechanism and supports targeted safety assessment.

For immune-mediated hepatitis, top enriched terms included PD-1 blockade immunotherapy, CD8⁺ T-cell downstream signaling, cytokine-cytokine receptor interaction, and TGF-β signaling pathways central to the pathogenesis of immune checkpoint inhibitors (i.e. CTLA-4 and PD-1/PD-L1) related liver toxicities. This is consistent with the strong concordance observed in mice, which are widely used to model T cell–mediated hepatic injury and immune-related AE *(Postow et al., 2018; Shojaie et al., 2021)*.

This integrated approach that links translatability metrics with mechanistic insights, provides a framework for identifying and prioritizing phenotypes with robust translational potential. These findings also help to support rational species selection, mechanistic interpretation and potential safety-target follow-up.

### Cross-Term Concordance Revealing Semantically and Mechanistically Related AEs

Building upon our identical-term mapping framework, we next sought to expand the translational landscape by systematically evaluating concordance among non-identical AE pairs. This comprehensive cross-term concordance analysis enabled detection of meaningful translational relationships that may be overlooked due to terminological differences between preclinical and clinical AE reporting. To maintain biological coherence, all cross-term associations were constrained within their respective SOC categories. Overall, this broader approach revealed 2,833 additional unique terms, including 2,375 clinical and 854 preclinical with 396 shared, from significant cross-term pairs. This expansion substantially broadened the spectrum of translational signals beyond the initial set captured through identical-term mapping. While stratified analyses by modality were conducted, the following results focus on the combined dataset across all modalities.

To quantitatively examine the relationship between semantic alignment and translational concordance within this expanded network of AE pairs, we integrated computational semantic similarity scoring. Embeddings for all MedDRA terms were generated using OpenAI text-embedding-3, and pairwise cosine similarities were calculated between preclinical-clinical AEs. These similarity scores were reported solely for descriptive and interpretive purposes, and were not applied as inclusion criteria, thresholds, or weighting factors in the statistical inference. As shown in **Fig. 6**, embedding-based similarity analysis revealed a statistically significant association between semantic similarity and translational strength as measured by LR+. Median LR+ values increased monotonically across semantic similarity quartiles, from 5.6 in the lowest quartile to 6.7 in the highest quartile (Kruskal–Wallis test, *p* = 1.8 × 10⁻¹⁰⁰). However, the broad and overlapping LR+ distributions within each quartile indicate that semantic similarity alone is insufficient to deterministically predict translatability or justify exclusion of low-similarity AE pairs. Several clinically important AEs exhibited strong predictive relationships with semantically similar preclinical phenotypes. For example, preclinical adrenal atrophy strongly predicted human adrenal suppression and adrenal insufficiency, conditions directly related to impaired adrenal function. Likewise, preclinical *Clostridioides difficile* colitis robustly anticipated human *C. difficile* infection, with the signal primarily driven by mouse studies. Examination of the associated drugs and targets indicates that this relationship was largely associated with broad-spectrum antibiotics acting on bacterial cell-wall synthesis pathways, including penicillin-binding proteins and β-lactamases. This observation is consistent with established mouse models of antibiotic-induced *C. difficile* colitis that recapitulate key features of the human disease, supporting the strong translational relevance of this phenotype (Chen et al., 2008; Best et al., 2012). Additional examples of high semantic similarity and strong concordance included preclinical eye inflammation predicting clinical eye irritation, and gastric ulcers robustly forecasting human gastrointestinal ulceration.

**Fig. 6.**
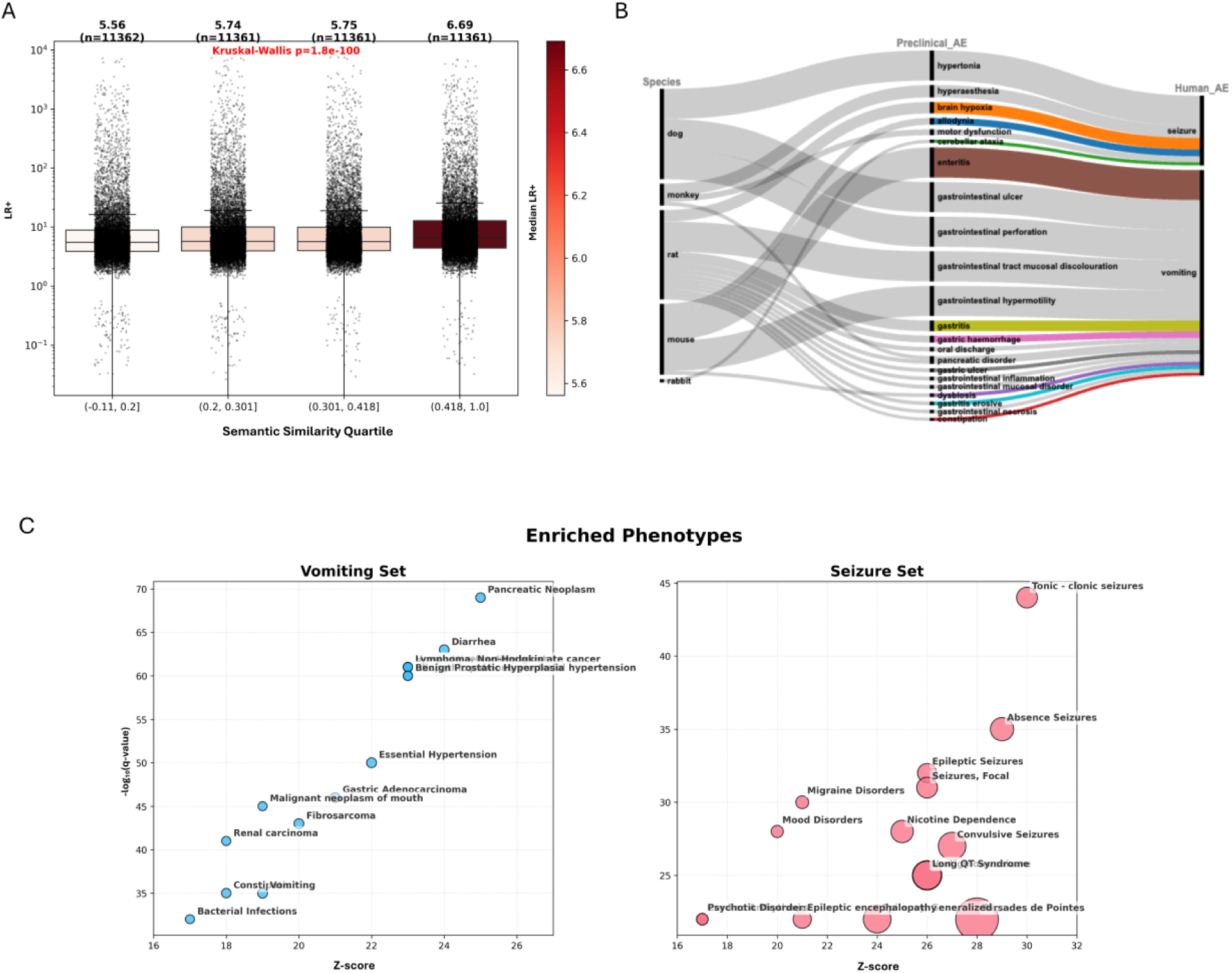
Cross-term AE concordance. (A) Distribution of LR+ across quartiles of semantic similarity for preclinical-clinical AE pairs. LR+ values and sample sizes are indicated above each box, with box colors scaled to the median value. (B) Sankey plot illustrating selected examples of significant, non-identical cross-species AE associations, with flow thickness proportional to LR+ value. (C) Bubble plots display enriched phenotypes among annotated targets and off-targets associated for drugs linked to vomiting (left) and seizure (right). Bubble size is proportional to the enrichment score.

Notably, a substantial proportion of adverse event pairs exhibited high translational concordance despite low semantic similarity, highlighting that meaningful preclinical-to-clinical translation can extend beyond surface-level terminological alignment and instead reflect deeper mechanistic or pathophysiological relationships. For example, the clinical endpoint “seizure,” had high translational concordance with multiple preclinical neurological phenotypes exhibiting low semantic similarity. These included hypertonia (LR+=∞, dog), hyperesthesia (LR+=40.7, monkey), brain hypoxia (LR+=38.1, rat), allodynia (LR+=22.7, mouse), motor dysfunction (LR+=19.6, monkey), and cerebellar ataxia (LR+=10.9, rabbit). Although these endpoints have limited direct lexical overlap with “seizure,” they represent drug-induced neural dysfunction or neurophysiological disturbances that can clinically manifest as seizures. On- and off-target enrichment analyses using the DisGeNET further supported these mechanistic links, revealing significant associations with seizure-related phenotypes (e.g., tonic-clonic seizures, absence seizures, focal seizures, and epileptic encephalopathy) and targets including voltage- and ligand-gated ion channels (e.g., CHRNA4/CHRNA7, GRIN2B, GRIA1), key receptors (DRD1–3, OPRK1, OPRM1) and neuromodulatory signaling modulators (HTR1A, SIGMAR1), all of which are implicated in seizure susceptibility and propagation (López-Meraz et al., 2005; Bozzi and Borrelli, 2013; Vavers et al., 2023; Zhang et al., 2024; Ferraro et al., 2025). A similar pattern emerged for the clinical AE “vomiting”, which demonstrate robust translational relationships with diverse preclinical gastrointestinal phenotypes, including enteritis and gastrointestinal hypermotility (LR+=∞, mouse), gastrointestinal ulcer and perforation (LR+=∞, dog), gastrointestinal mucosal discoloration (LR+=∞, rat), gastritis (LR+=35.6, rat), gastric hemorrhage (LR+=22.6, rat), and pancreatic disorders (LR+≈12–14, rat and monkey). While these preclinical terms are not direct synonyms for “vomiting,” they reflect underlying gastrointestinal injury, inflammation, and motility disturbances central to emetic biology. Enrichment analysis highlighted targets and pathways involving serotonergic signaling (HTR3A-D), opioid receptors (OPRM1), gut–brain axis regulators (GLP1R, GDF15), and excitability-related channels (CACNA1A), providing mechanistic support for these translational links (Niesler et al., 2008; Joy Lin et al., 2014; Sugino et al., 2014; Borner et al., 2020; Indelicato and Boesch, 2021; Huang et al., 2024). Together, these findings demonstrate how incorporation of pharmacological target information enables a more mechanistically grounded assessment of translational relevance.

Overall, these results establish that extending beyond direct term matches to include semantically and mechanistically related AE pairs substantially expands the scope of detectable translational signals between preclinical and clinical data. This expanded mapping framework uncovers both expected and previously unrecognized associations, providing a richer, more nuanced basis for evaluating the biological relevance of preclinical findings and informing risk assessment.

### Integrated Web Application and Multi-Agent AI System

To maximize the impact and accessibility of our concordance resource, we developed an integrated suite consisting of a user-friendly web application. This platform is designed to democratize access to our large-scale concordance results and to enable users to search for AE translatability scores categorized by different modalities and species, facilitating the evaluation of the translatability of findings from animal models to humans and supporting informed preclinical species selection. To elucidate toxicity mechanisms, the app offers two main search functionalities: (1) querying drugs and their on- and off-targets associated with specific AEs, and (2) identifying AEs linked to particular drugs or targets, grouped by species (**Supplementary Figure 4**).

Anticipating the future needs by leveraging the increasingly powerful agentic artificial intelligence, we further explored how our concordance data can be integrated into real-world decision-making scenarios facilitated by AI (**Fig. 7**). To this end, we developed a multi-agent AI system ToxAgents, consisting of two agents: an investigative toxicology scientist agent and a regulatory affairs expert agent, representing the scientific research and regulatory sides, respectively, of an investigative toxicology group in a pharmaceutical company. The investigative toxicity scientist agent can utilize two tools: concordance data (discussed above) and a safetyome tool, a curated resource centered on drug safety–critical genes/proteins and their associated AEs. The foundational approach for the safetyome was described previously *(Liu et al., 2021)*, and has since been further enhanced in our research. In parallel, the regulatory affairs expert agent brings specialized knowledge of ICH safety guidelines. Within the web interface, users can manually query concordance metrics and associated targets, while ToxAgents extends this functionality by providing a standardized, evidence-integrated interface to the concordance statistics. This supports species-to-human translatability assessment, quantitative risk inference, and mechanistic hypotheses generation aligned with ICH safety assessment principles. To demonstrate the capabilities of our multi-agent AI system, ToxAgents, we present three case studies comparing results from ToxAgents with native responses generated by GPT-4.1. These scenarios are to simulate some of the common challenges faced during day-to-day drug development: 1. An adverse event is observed in a preclinical toxicology species, and we hope to determine how translatable the phenotype is to humans and what potential mechanisms could be investigated. 2. We have a phenotype of interest and need to identify the most appropriate species for testing this AE in a preclinical setting. 3.We have a human AE that is known to occur with other drugs or within the competitive landscape, but is not well reproduced in a particular species, and we need to determine alternative endpoints to monitor that may translate to the AE of interest in humans. Additional illustrative examples are available within the ToxAgents package.

**Fig. 7.**
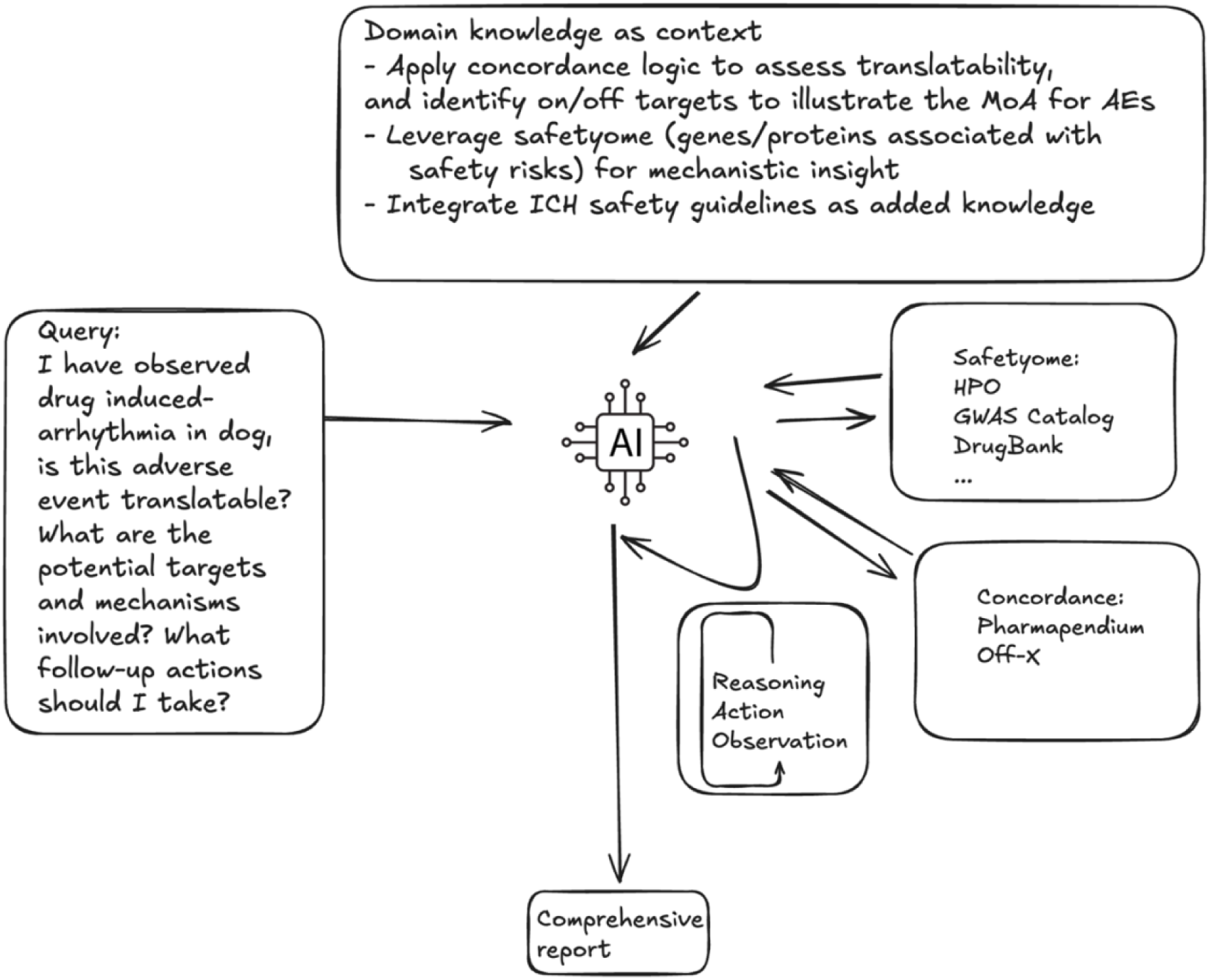
AI-enabled framework for evaluating the translatability of preclinical AEs. The process integrates domain knowledge, including concordance logic and data schemas, mechanistic insights from on/off targets and safetyome, and ICH safety guidelines. The AI system synthesized these inputs through reasoning, and observation-action loops to generate a comprehensive, context-aware, data-driven report with actional recommendations for follow-up.

*Case Study 1 – I have observed drug induced-arrhythmia in dog, is this adverse event translatable? What are the potential targets and mechanisms involved? What follow-up actions should I take?*

Both approaches recognize that arrhythmias observed in dogs are highly translatable to humans. With respect to underlying mechanisms, both analyses highlight key genetic contributors, such as the KCNH2 gene and cardiac ion channels. However, the ToxAgents excels by providing the quantified positive likelihood ratio (LR+) for dogs, estimated as 3.48 × 10⁹, demonstrating strong confidence in the translatability. Importantly, ToxAgents provides ranked, evidence-weighted prioritization of targets, such as KCNQ1, SCN5A, KCND3, and KCNA5 among others, for follow-up mechanistic investigations, which represents an added layer not produced by native LLM output alone. In addition, the system incorporates regulatory-relevant parameters such as the hERG IC₅₀/Cmax safety margins, providing operational context for cardiac-safety evaluation. Collectively, while native LLM responses remain broad and qualitative, ToxAgents delivers reproducible, drug-derived quantitative metrics, mechanistic prioritization, and regulatory-aligned guidance that are directly grounded in the concordance framework presented here.

*Case Study 2 – Which species would be most suitable for preclinical studies on hematuria? What are the potential targets and mechanisms associated with hematuria findings in selected preclinical species? If I observe adverse effects related to hematuria, what follow-up actions should I consider?*

Both approaches agree on the importance of dogs as preclinical models for evaluating hematuria. Both analyses discuss similar mechanistic pathways, such as glomerular and tubular injury, vascular disruption, and coagulation disturbances and recommended follow-up actions. However, there’s a divergence between the analyses concerning the predictive value of rodents. The native ChatGPT response indicates that rodents, particularly rats, are frequently chosen due to practicality, and extensive historical use, but might have less predictivity. In contrast, the ToxAgents identified mice and rats as suitable preclinical species based on quantitative concordance data. It reports robust LR+ (21.1 for mice and 10.3 for rats), suggesting strong predictive translatability to humans. The native ChatGPT analysis primarily relies on general literature and traditional understanding and knowledge, whereas the ToxAgents draws directly from the concordance data with quantitative metrics, providing a data-driven perspective. Additionally, ToxAgents explicitly identifies particular targets, notably COL4A3 and COL4A4, which encode type IV collagen essential to the integrity of the glomerular basement membrane and mechanistically link to hematuria. This strengthens mechanistic insights and facilitates targeted experimental follow-up, which contrasts with native ChatGPT’s broader discussion lacking detailed mechanistic connections. Furthermore, ToxAgents delivers clearly structured, actionable follow-up recommendations aligned explicitly with regulatory frameworks. These include defined operational thresholds, such as recommended safety margins (≥10-fold exposure margin according to ICH M3(R2)). In contrast, the native ChatGPT response, while practically relevant, does not clearly specify detailed regulatory criteria, somewhat limiting its immediate translational utility. In summary, ToxAgents provides explicit quantitative evidence, precise mechanistic detail, and structured regulatory guidance, compared to the broader, literature-based reasoning provided by native ChatGPT.

*Case Study 3 – I want to know what adverse effects in rat other than vomiting best translate into vomiting in humans for small molecule drugs. also, what’s the potential biological mechanism causing the adverse effect*.

Both the native ChatGPT and ToxAgents analyses recognize the fundamental challenge that rats are unable to vomit. ChatGPT highlights “pica” (kaolin consumption) as the most widely used behavioral marker in rats for nausea and vomiting, and noting other nonspecific indicators such as hypoactivity, salivation, and piloerection. The ToxAgents systematically ranks the most predictive surrogate adverse effects in rats based on LR+ from the concordance analysis, providing robust quantitative support for their translational relevance to human vomiting. For example, it identifies gastritis (LR+ = 29.86), pancreatic disorder (LR+ = 16.95), erosive gastritis (LR+ = 15.55), gastric ulcer (LR+ = 13.68), and oral discharge (LR+ = 11.97) as the most predictive non-vomiting rat adverse effects for human emesis, far exceeding the qualitative evidence provided by the ChatGPT analysis. In addition, the ToxAgents offers detailed mechanistic insights by specifying the key molecular targets and pathways associated with each adverse effect. The analysis cites serotonin 5-HT3 receptors, dopamine D2 receptors, neurokinin-1 receptors, muscarinic and histaminergic receptors, prostaglandin pathways, and others as being repeatedly implicated in both preclinical and clinical settings. This level of detail enables a mechanistic linkage between observed preclinical effects and clinical outcomes, enhancing the interpretability of preclinical data. By comparison, while the native ChatGPT analysis is comprehensive in describing surrogate behaviors, mechanisms, and the broader context, it remains primarily qualitative and lacks the structured and quantitative approach. Therefore, ToxAgents delivers superior empirical rigor, quantitative evidence, and explicit mechanistic detail, making it more robust and practical for real-world application.

For these case studies, we also assessed reproducibility and robustness as described in the Methods section. Across all three scenarios, LR scores and species were consistently and correctly reported, demonstrating that the agents can accurately query the database to retrieve required information (**Supplementary Table 8 & 9**). This outcome was expected, as the system had already been thoroughly validated during tool testing to ensure correct query and database access. The full-answer semantic similarity is above 0.9 across all cases. Target/mechanism identification achieved similarity scores of 0.92 for arrhythmia in dog, and 0.81 and 0.74 for hematuria and vomiting, respectively. These lower scores for hematuria and vomiting likely reflect the greater complexity of underlying mechanisms, so that even with prioritization based on target-phenotype associations from safetyome, some variability remained in the generated reports. A similar pattern was observed for follow-up action recommendations. In the robustness analysis, robustness decreases minimally across cases, indicating stable outputs under prompt variation. Overall, ToxAgents produced consistent, high-similarity outputs across repeated runs and maintained stability when queries were rephrased, supporting its reliability for translational safety interrogation.

In summary, this integrated infrastructure not only enhances transparency and user engagement but also fosters a culture of open science. By making concordance analytics and mechanistic insights available, our web application and multi-agent AI system provide a robust, systematic, scalable and much needed platform to offer translational insights in safety assessment, facilitate regulatory interactions, and ultimately guide more effective preclinical tests as well as support the development of safer, more effective therapeutics for all.

## CONCLUSIONS

By assembling and harmonizing a large concordance dataset spanning both marketed and investigational drugs across multiple species, this study aims to make a constructive contribution to translational safety science.

At the organ system level, concordance was generally low to modest. Positive predictivity was strongest for infections and infestations, followed by immune, renal-urinary, and musculoskeletal disorders, whereas general disorders, nervous system, and pregnancy/perinatal conditions ranked lowest. Biologics exhibited higher concordance for immune-related disorders than small molecules, and monkeys showed higher median predictivity than other species, particularly for infections, immune, and gastrointestinal disorders. Negative predictivity was generally low, indicating limited ability to exclude human AEs based on their absence in animals. At the phenotype level, concordance signals became more specific and decision relevant. We identified 850 significant identical-term events, including 780 for small molecules, 237 for biologics, and 94 within the exposure-controlled subset. Extending beyond identical-term mapping, inclusion of semantically related and mechanistically related AE pairs uncovered 2,833 additional unique endpoints. Integrating both on- and off-target annotation strengthened mechanistic interpretation by highlighting biological pathways and potential safety targets for subsequent validation.

To ensure that these advances translate into day-to-day impact, we developed the web app and the multi-agent AI system (ToxAgents), which transform concordance analytics into actionable decision support. The app provides an interactive interface for exploring concordance results, and the ToxAgents layers on structured, role-aligned reasoning for translational safety. The investigative scientist agent interrogates the concordance data, together with the safetyome of safety-critical targets, to assemble mechanistic rationales and prioritize candidates for further evaluation. In parallel, the regulatory-affairs agent interprets outputs in the context of ICH safety guidelines and organized recommendations for study design. Together, the agents synthesize search results to support human risk assessment, species selection, mechanistic follow-up, and preparation for regulatory interactions. This architecture facilitates broad, standardized access to concordance analytics, promoting wider adoption across scientific and pharmaceutical communities.

Despite these advances, several challenges remain inherent to the landscape of translational safety assessment. A primary limitation relates to the subjectivity and granularity of source data. While OFF-X and PharmaPendium offer unparalleled breadth and regulatory rigor, the assignment of MedDRA terms, particularly for preclinical findings such as animal behaviors or nuanced histopathological changes, may involve subjective interpretation by original authors or database curators *(Clark and Steger-Hartmann, 2018)*. In some cases, this can lead to ambiguity or inconsistency in how observations are classified and mapped, especially for endpoints not directly translatable between species. Additionally, missing AEs in approval packages or database sources due to incomplete or selective reporting, or failure to extract relevant findings can potentially mask genuine translational signals. Another limitation is that although OFF-X includes investigational drugs with daily updates, reported AEs may be neutral or lack severity because companies may choose not to disclose more impactful safety findings. Moreover, the present analysis encodes AEs as binary (present/absent) outcomes for each drug, without quantitative metrics on frequency, severity, or explicit dose-response relationships. This reflects the nature of reporting within the source databases, where comprehensive incidence data and dosing details are not consistently available. As such, while our findings are robust for mapping qualitative translational signals and identifying mechanistic links, they are less suited for assessing clinical risk thresholds or for guiding nuanced decision-making where quantitative data would be essential. Mechanistically, our framework harnesses drug target annotation to illuminate the biological processes underlying concordant AEs in human and preclinical species. Nonetheless, this approach can be confounded by the complex pharmacology of many drugs, which often engage multiple on- and off-targets—some unrelated to the observed toxicities. Moreover, adverse effects may result from metabolites or from off-target interactions that are not yet characterized.

Looking ahead, future methodological refinements should leverage increasingly available quantitative data on incidence, severity, and dose-response relationships to enhance translational safety assessment. Stratifying drugs by therapeutic indication such as oncology versus non-oncology indications, may further enable targeted and clinically relevant predictions. Integrating advanced computational approaches, including network-based and machine learning approaches, can reveal higher-order translational patterns and multi-event associations, offering deeper insight into the complex, interconnected nature of clinical toxicities. Continued advancement of multi-agent, AI-enabled platforms with iterative refinement and expanded mechanistic tools such as multi-omics integration and automated pathway inference will support broader adoption and foster cross-sector collaboration. A dedicated pipeline will facilitate timely retrieval and analysis, allowing concordance metrics to be refreshed regularly and ensuring that the framework evolves with the expanding volume of curated safety information.

In summary, the integrative concordance framework presented here establishes a scalable and accessible foundation for advancing translational safety science. By addressing key challenges in event mapping, stratification, and accessibility, this study sets the stage for more actionable and mechanism-informed risk assessment, with the ultimate goal of improving therapeutic safety and patient outcomes.

